# Matrix vesicle isolation from a three-dimensional *in vitro* bone model

**DOI:** 10.1101/2023.01.10.523451

**Authors:** Sana Ansari, Lotte van Dasler, Yuana Yuana, Miguel Castilho, Keita Ito, Sandra Hofmann

**Author notes:** Corresponding author: Sandra Hofmann.

## Abstract

Extracellular vesicles (EV) are nano-sized bilayer vesicles that are involved in biological functions and secreted by a wide variety of cells. Osteoblasts, the bone forming cells, can release a subset of EVs known as matrix vesicles (MtVs) which are believed to be involved in matrix mineralization and feature bone forming properties. Osteoblast-derived EVs or MtVs have been mostly isolated from conditions which are still far from nature, i.e. mesenchymal stromal cells (MSCs), or osteoblast cell lines cultured in two-dimensional (2D) tissue culture flasks. In our study, we aimed at investigating whether MtVs could also be isolated from an environment which better resembles the complex *in vivo* situation. This study investigated the EVs secretion during osteogenic differentiation of human bone marrow MSCs (hBMSCs) in the most advanced human three-dimensional (3D) *in vitro* woven bone constructs previously developed by our group. hBMSCs were cultured in spinner flask bioreactors which induced wall shear stress on cells and directed the cells to differentiate towards osteoblasts and osteocytes. The EVs secreted into the culture medium were isolated and characterized based on their morphological, biological, and functional properties. The characteristics of a part of isolated EVs shared similarities with MtVs. These vesicles were electron-dense and electron-lucent, showed alkaline phosphatase (ALP) activity, increased the amount of released free phosphate into the culture medium, and increased the amount of deposited phosphate within the ECM. The results indicate that a complex 3D environment mimicking bone development is favorable to stimulate MtV-producing cells to produce targeted MtVs *in vitro*. These MtVs potentially could be used as a biological agent for bone regeneration and fracture healing through, for instance, integration with biomaterials to target bone formation locally.

## 1- Introduction

Bone is a rigid connective tissue that provides structural support for the body, allows movement, protects internal organs, and serves as storage for calcium and growth factor. Despite its passive appearance, bone is a highly dynamic tissue that continuously undergoes a physiological process called bone remodeling [1]. This process maintains bone strength and mineral homeostasis through close interaction between bone-forming osteoblasts, bone-resorbing osteoclasts, and bone-regulating osteocytes [1]. Osteoblasts are bone forming cells derived from mesenchymal stromal cells (MSCs). Under appropriate mechanical and/or chemical stimuli, MSCs differentiate towards osteoblasts [2]. Osteoblasts are responsible for the formation of the bone mineralized tissue which consists of mainly collagen type 1 and carbonated hydroxyapatite deposited within/outside of these collagen fibrils, forming a continuous inter-connected cross-fibrillar pattern [3]. This composite matrix is highly organized at multiple hierarchical levels giving bone its mechanical properties [4].

Coordinating the regulatory processes between bone specific cells during bone remodeling is thought to happen partly through extracellular vesicles (EVs) produced by each cell type [5]. EVs have complex membrane structures and can be secreted by almost all cell types, including osteoblasts. EVs are phospholipid-enclosed nanoparticles containing lipids (to protect the bioactive substances within EVs), proteins (to give them targeting abilities), and nucleic acids (to transmit information to other cells) [5]. Osteoblast-derived EVs have been shown to be involved in multiple processes such as osteogenic differentiation, osteoclast formation, and inorganic matrix deposition [6–9]. A subset of osteoblast-derived EVs is known as matrix vesicles (MtVs). These vesicles range in size from 50-1000 nm and can bind to the collagenous matrix [10]. The membrane of these vesicles is enriched with phosphatidylserine (PS)-binding annexin proteins such as annexin A5 and phosphatases such as alkaline phosphatase (ALP). These membrane proteins facilitate entry of calcium and phosphate ions into MtVs. These ions then form amorphous or crystalized minerals inside the MtVs which can rupture the membrane and form mineral nodules in the extracellular matrix (ECM) [10,11]. MtVs have shown to target bone *in vivo* and induce bone formation *in vitro* and *in vivo* [12–15]. For instance, EVs derived from the osteoblast cell line MC3T3-E3 added to bone marrow derived mesenchymal stromal cells (BMSCs) accelerated the osteogenic differentiation of BMSCs and mineral deposition [12,13]. Nevertheless, these osteoblast-derived EVs or MtVs studies have been mostly isolated from cells cultured in two-dimensional (2D) tissue culture flasks. In our current study, we investigated whether MtVs could be isolated from cells cultured in the 3D human *in vitro* woven bone constructs, an environment with more similarity to the complex environment of *in vivo* situation.

We have previously developed a functional 3D self-organizing co-culture of osteoblasts and osteocytes that represents the woven bone formation using human bone marrow MSCs (hBMSCs) [16]. Woven bone forms during skeletal development where rapid pace of matrix mineralization is needed [17]. In this process, pre-osteoblasts lay down randomly oriented collagen that becomes highly mineralized [18]. The formation of highly mineralized matrix in this 3D *in vitro* woven bone construct could be partly associated to the release of MtVs by osteoblasts. In our current study, we used this setup and isolated the osteoblast-derived EVs or MtVs released by cells in the cell culture supernatant. We hypothesized that those EVs would show similar characteristics as MtVs and promote collagenous matrix mineralization *in vitro*.

To create the woven bone constructs, hBMSCs were seeded on silk fibroin scaffolds. These cell-seeded constructs were placed inside a spinner flask bioreactor which induce continuous wall shear stress on cells for 4 weeks (Figure 1A). After concentrating cell culture supernatant, EVs were isolated using size-exclusion chromatography (Figure 1B) and characterized based on their morphological, biological (Figure 1C), and functional properties (Figure 1D). The culture medium for hBMSCs differentiation contained fetal bovine serum (FBS). To account for EVs that were already present in FBS, EVs from FBS containing medium were also isolated and characterized using the same approach. To study the potential of isolated EVs on osteogenic differentiation, the EVs from cell culture supernatant (Cell-EVs) were added to hBMSCs cultured in the previously developed serum substitute medium (SSM) for 3 weeks [19]. The influence of MtVs on osteoblast differentiation of hBMSCs and deposition of mineralized matrix were studied.

**Figure 1.**
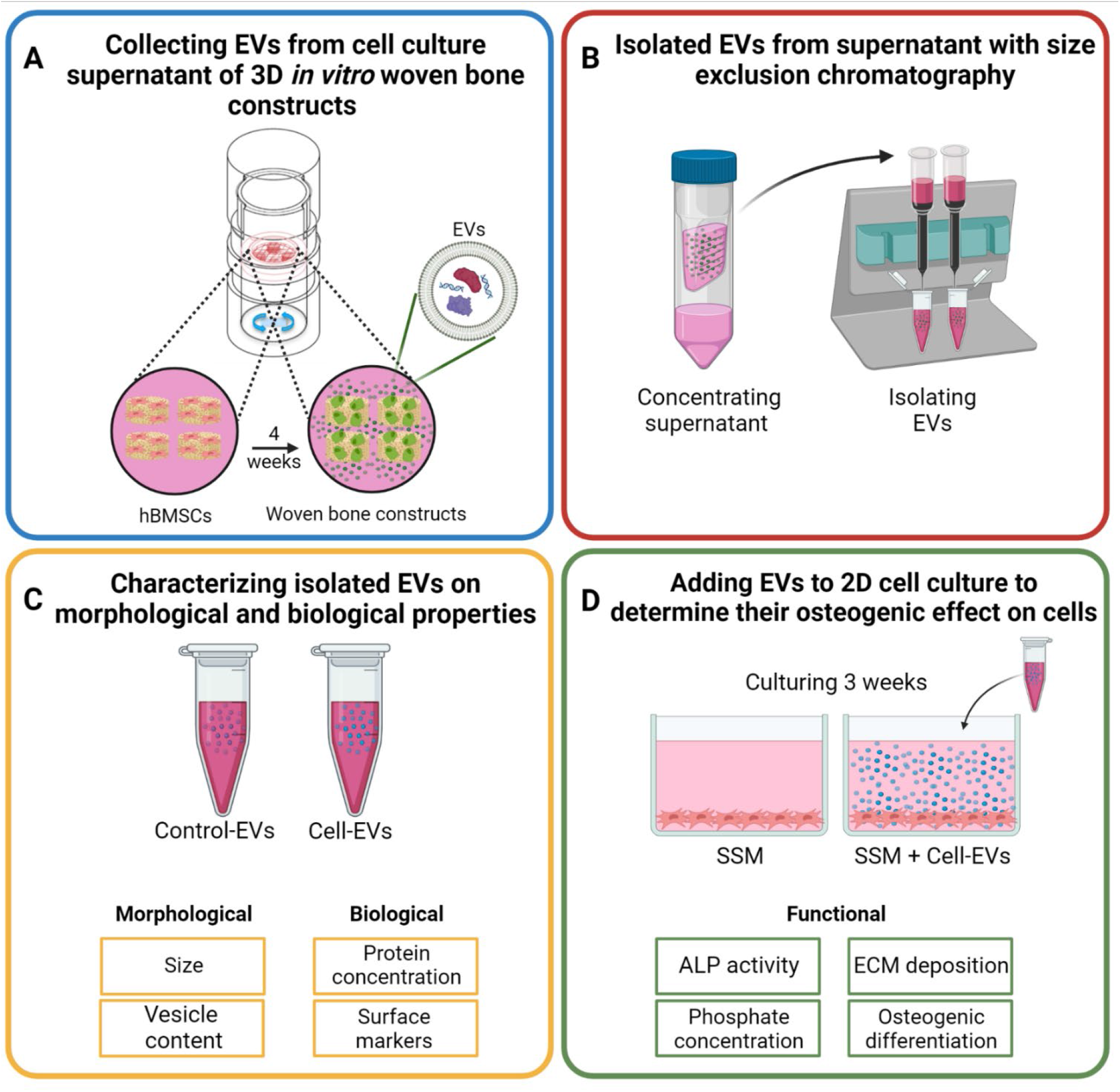
*Isolation and characterization of MtVs from cell culture supernatant of osteogenic differentiated hBMSCs during in vitro* woven bone formation. hBMSCs were cultured under continuous shear stress for 4 weeks which could release EVs to the medium (A). Cell culture supernatant was first concentrated using a spin-column and then EVs were isolated from the collected cell culture supernatant (Cell-EVs) and control group (Control-EVs) using size-exclusion chromatography (B). Isolated EVs were characterized based on their morphological and biological properties (C). hBMSCs were cultured in serum substitute medium (SSM) containing Cell-EVs for 3 weeks to study the influence of EVs on the osteogenic differentiation process (D). Created with BioRender.com.

## 2- Material and Methods

### 2-1 Materials

Dulbecco’s modified eagle medium (DMEM high glucose, 41966 and low glucose, 22320), non-essential amino acids (NEAA, 11140050), antibiotic/antimycotic (Anti-Anti, 15240062), and trypsin-EDTA (0.5%, 2530054), Insulin-Transferrin-Selenium (ITS-G, 41400045), GlutaMAX (35050061), chemically defined lipid concentrate (CD-lipid, 11905031) were from Life Technologies (The Netherlands). For cell expansion, FBS (F7524) from Sigma-Aldrich (The Netherlands) and for osteogenic differentiation, FBS (SFBS) from Bovogen (Australia) were used. Basic fibroblast growth factor (b-FGF, 100-18B) was purchased from Peprotech (UK). Recombinant human bone morphogenetic protein-2 (rhBMP-2, 7510200) was purchased from Medtronic Sofamor Danek (USA). Human serum albumin (HSA, A1653) was purchased from Sigma-Aldrich (The Netherlands). Unless noted otherwise, all other substances were of analytical or pharmaceutical grade and obtained from Sigma-Aldrich (The Netherlands).

### 2-2 Silk fibroin scaffold fabrication

Silk fibroin scaffolds were prepared by cutting and cleaning 3.2 grams of Bombyx mori L. silkworm cocoons (Tajima Shoki., Ltd. Japan). The cocoons were then degummed by boiling in 1.5 L ultra-pure water (UPW) containing 0.02 M Na_2_CO_3_ (Sigma-Aldrich, S7795) for 1 hour, whereafter it was rinsed with 10 L cold UPW to extract sericin. Dried purified silk fibroin was dissolved in 9 M lithium bromide (LiBr, Acros organics, 199870025) solution in UPW at 55°C for 1 hour and dialyzed against UPW for 36 hours using SnakeSkin Dialysis tubing (molecular weight cut-off (MWCO): 3.5 kDa, Thermo Fisher Scientific, The Netherlands). The silk fibroin solution was frozen at -80°C for at least 2 hours and lyophilized (Freezone 2.5, Labconco, USA) for 4 days. 1.7 grams of lyophilized silk fibroin was then dissolved in 10 mL 1,1,1,3,3,3-Hexafluoro-2-propanol (HFIP, Fluorochem, 003409) at room temperature for 5 hours resulting in a 17% (w/v) solution. One mL of silk-HFIP solution was added to a Teflon container containing 2.5 grams NaCl with a granule size between 250-300 μm. After 3 hours, HFIP was allowed to evaporate for 4 days in a fume hood. Silk fibroin-NaCl blocks were immersed in 90% (v/v) methanol (Merck, The Netherlands) in UPW for 30 minutes to induce the protein conformational transition to β-sheets and let dry overnight [20]. Scaffolds were cut into disks of 3 mm height with an Accutom-5 (Struer, Type 04946133, Ser. No. 4945193), followed by immersion in UPW for 2 days to extract NaCl. Disc-shaped scaffolds were made with a 5 mm diameter biopsy punch (KAI medical, Japan) and autoclaved in phosphate buffered saline (PBS, Sigma-Aldrich, P4417) at 121°C for 20 minutes.

### 2-3 Medium collection and EV isolation

#### 2-3-1 Cell expansion and seeding

The hBMSCs were isolated from human bone marrow (Lonza, USA) and characterized as previously described [21]. Passage 4 hBMSCs were expanded (2500 cell/cm^2^) in expansion medium (DMEM high glucose containing 10% FBS Sigma, 1% Anti-Anti, 1% NEAA, and 1 ng/ml b-FGF) for 9 days, the medium was replaced 3 times per week. At day 9, cells were 80% confluent and trypsinized using trypsin-EDTA and dynamically seeded on scaffolds as previously described [22]. Briefly, each scaffold (n=8) was incubated with a cell suspension (1*10^6^ cells/4 mL control medium, i.e. DMEM, 10% FBS Bovogen, and 1% Anti-Anti) in 50 mL tubes placed on an orbital shaker at 150 rpm for 6 hours in an incubator at 37°C. Then, cell-seeded scaffolds were transferred to spinner flask bioreactors (n=4 per bioreactor). Each bioreactor contained a magnetic stir bar and was placed on a magnetic stirrer (RTv5, IKA, Germany) at 300 rpm in an incubator (37°C, 5% CO_2_) [23]. Each bioreactor was filled with 5 mL control medium containing osteogenic differentiation factors (50 μg/mL ascorbic-acid-2-phosphate, 100 nM dexamethasone, and 10 mM β-glycerophosphate, β-GP). The medium was refreshed 3 days a week for 4 weeks.

#### 2-3-2 Extracellular vesicle (EV) isolation

Cell culture supernatant (5 mL from each bioreactor) was collected at each medium change and centrifuged at 200g for 10 minutes. The supernatant was transferred to a new tube and centrifuged again at 2000g for 10 minutes to remove dead cells and debris. The supernatant was collected in a new tube, frozen in liquid nitrogen, and stored at -80°C. This process was done after each medium change for 4 weeks of osteogenic differentiation of hBMSCs in spinner flask bioreactors. To isolate EVs, the stored medium aliquots were thawed at 37°C and combined. Fifteen mL of the combined medium was concentrated using Amicon Filter Units (MWCO=100 kDa, ACS510024, Merck Millipore) centrifuged at 4000g for 15 minutes. EV isolation was done using qEV1/70 nm column (IC1-70, Izon Science, France). As mentioned in the supplier’s user manual, these columns are suitable to isolate vesicles in the range of 70-1000 nm. The columns were prepared before use as described in the user manual. Briefly, the columns were rinsed with 11.5 mL 0.5M filtered NaOH which was followed by 2 times rinsing with 11.5 mL filtered UPW. Then, the columns were rinsed with 11.5 mL 20% ethanol and 2 times with 11.5 mL sterile PBS. The concentrated medium (around 1 mL) was overlaid on the column. The sample was allowed to run completely into the loading frit, the layer separating loading reservoir from the gel part of the column. Once the sample has passed into the frit, 3 mL filtered PBS was added to the column. Four mL of the flowthrough PBS was discarded as there should be no particles in this volume. We added 2.8 mL of filtered PBS again and collected 2.8 mL of this flowthrough directly. This collected fraction should contain the EVs. The collected EVs were concentrated again using Amicon Filter Units resulting in up to 500 μL of EV solution. These EVs were referred as Cell-EVs in this study. The culture medium for hBMSCs differentiation contained FBS. To account for EVs that were already present in FBS, the same procedure was done on 15 mL of control medium (DMEM, 10% FBS Bovogen, and 1% Anti-Anti). EVs isolated from control medium were referred as Control-EVs. The EVs were characterized and used in the cell culture experiment as follows.

### 2-4 Characterization of EVs

#### 2-4-1 Bicinchoninic acid (BCA) assay

Protein content of EVs was determined using Micro BCA protein assay kit (23235, Thermo Fisher Scientific, The Netherlands) according to the manufacturer’s instructions. Briefly, 150 μL of EV solution was mixed with 150 μL of Micro BCA working Reagent in a 96-well plate. The plate was covered with a sealing tape and incubated at 37°C for 2 hours. Absorbance was measured at 562 nm on a plate reader and protein concentration was calculated by comparison to standards of known bovine serum albumin (BSA) concentrations.

#### 2-4-2 Alkaline phosphatase (ALP) assay

The membrane of MtVs is equipped with mineralization-specific components such as ALP. To determine whether the isolated EVs from the cell culture supernatant contained active ALP, the ALP activity of EVs was measured as follow. EV solution (240 μL) was mixed with 240 μL of 0.2% (v/v) Triton X-100 and 5 mM MgCl_2_ solution and incubated for 30 minutes at room temperature. In a 96-well plate, 80 μL of the samples was mixed with 20 μL of 0.75 M 2-amino-2-methyl-1-propanol buffer and 100 μL of 10 mM p-nitrophenylphosphate solution and incubated until colour developed, before 0.2 M NaOH were added to stop the conversion of p-nitrophenylphosphate to p-nitrophenol. Absorbance was measured in at 405 nm on a plate reader and ALP activity was calculated by comparison to standards of known p-nitrophenol concentration.

#### 2-4-3 Measure the amount of free phosphate in medium with/without the presence of EVs

To determine the potential of EVs on increasing the concentration of phosphate in the medium, the serum substitute medium was supplemented with EVs and 10 mM β-GP. β-GP is a phosphate source in osteogenic differentiation cultures that can be cleaved by ALP (expressed by osteoblasts during differentiation or present in FBS [24]) and released free phosphate to the medium. Since the membrane of MtVs are equipped with ALP, it was hypothesized that isolated EVs can increase the amount of the free phosphate in the medium containing β-GP. To measure the amount of free phosphate in the medium, serum substitute medium with and without EV supplementation (n=3 per group) was incubated with β-GP at 37°C for 48 hours. The phosphate concentration was then measured according to the manufacturer’s instructions (Malachite Green Phosphate Assay Kit, Sigma-Aldrich, The Netherlands). Briefly, 80 μL of 1:200 (v/v) diluted samples in UPW were mixed with 20 μL of working reagent and incubated for 30 minutes at room temperature. In this assay, a green complex is formed between molybdate and free phosphate. Absorbance was measured in at 620 nm on a plate reader and phosphate concentration was calculated by comparison to a phosphate standard provided in the kit.

#### 2-4-4 Nanoparticle tracking analysis (NTA)

NTA was used to determine the size and concentration of EVs. This method is based on the Brownian motion of individual particles in solution. The movement of the particles is tracked based on light scattering. A Nanosight NS300 instrument (Malvern Panalytical) equipped with a scientific complementary metal-oxide-semiconductor (sCMOS) camera was used. The camera was mounted on an optical microscope, allowing visualization of the light scattered by the injected particles that were present in the focus of an 80 μm beam generated by a single mode laser diode with a blue laser (488 nm). During the measurement, temperature was kept at 24°C. The samples were diluted 10 times in filtered PBS and 1 mL of the samples was injected into the Nanosight chamber. One mL filtered PBS was used as a control. NTA 3.2 software was used to analyze the data. Five videos of 60 seconds were captured per sample at camera level 10 and screen gain of 5. During the data analysis the screen gain was set to 5 with the detection threshold of 10.

#### 2-4-5 Flow cytometry

Flow cytometry was used to detect and measure the proportion of Annexin A5 and CD-9, as typical EV surface markers. Briefly, 20 μL of EV solution was first mixed with 20 μL of flow cytometry buffer, i.e. 1% BSA dissolved in PBS containing 1% binding buffer (521103674, Miltenyl Biotec). The binding buffer contains calcium which is needed for the binding of Annexin A5 to phosphatidylserine (PS). Next, either 1 μL of Annexin A5 conjugated with FITC (BioLegend, 640906) and/or 1 μL of CD-9 antibodies conjugated with APC (BioLegend, 312107) was added to the samples. The samples were then incubated at 4°C for 30 minutes in the dark. Another 260 μL flow cytometry buffer was added to each sample prior to the measurement done by FACSymphony A3 (660937, BD Biosciences). The triggering threshold was set on forward scatter at 200. All samples were analyzed at low speed (35 μL/second) for 2 minutes. The obtained results were analyzed with FlowJo (v10.8.1, BD Biosciences).

#### 2-4-6 Cryo-Transmission electron microscopy (TEM)

Vitrified thin films for CryoTEM analysis were prepared using an automated vitrification robot (FEI Vitrobot Mark IV) by plunge vitrification in liquid ethane. Before vitrification, a 200-mesh copper grid covered with a Quantifoil R 2/2 holey carbon film (Quantifoil Micro Tools GmbH) was surface plasma treated for 40 seconds using a Cressington 208 carbon coater. CryoTEM imaging was carried out on the cryoTITAN (Thermo Fisher, previously FEI), equipped with a field emission gun (FEG), a post-column Gatan imaging filter (model 2002) and a post-GIF 2k × 2k Gatan CCD camera (model 794). The microscope was operated at 300 kV acceleration voltage in bright-field TEM mode with zero-loss energy filtering at a nominal magnification of 6.500×; or at 24.000× magnification both with a 1s image acquisition time.

### 2-5 Addition of EVs to osteogenically differentiating hBMSCs

To determine the influence of EVs on osteogenic differentiation, hBMSCs were seeded on well-plates and cultured in the previously developed serum substitute medium (SSM) [19]. The use of serum substitute medium was to avoid the influence of EVs present in FBS [25].

#### 2-5-1 Cell expansion and seeding

Passage 4 hBMSCs were expanded (2500 cell/cm^2^) in expansion medium (DMEM high glucose containing 10% FBS Sigma, 1% Anti-Anti, 1% NEAA, and 1 ng/ml b-FGF) for 9 days, the medium was replaced 3 times per week. At day 9, cells were 80% confluent, trypsinized and seeded on vitronectin coated 48-well plate (Greiner bio-one, CELLSTAR, 677-180). The wells of a 48-well-plate were coated with 5 μg/ml vitronectin (Peprotech, 140-09) diluted in PBS to increase cell attachment to the surface of well plate in the serum substitute medium [19]. In short, 100 μL of vitronectin solution was added to each well and incubated in an incubator (37°C, 5% CO_2_) for 2 hours and then at 4°C overnight. The following day, the vitronectin solution was aspirated, the wells were rinsed 3 times with PBS, and the well-plate was pre-warmed in an incubator before seeding cells into the wells. We seeded 2500 cells per well resulting in 2500 cells/cm^2^ and cultured them in serum substitute medium (Table 1), additionally supplemented with osteogenic differentiation factors (50 μg/mL ascorbic-acid-2-phosphate, 100 nM dexamethasone, 10 mM β-glycerophosphate). The medium was refreshed 3 days a week during the first week. After 1 week of culture, a group of cells was continued to be cultured without EVs in serum substitute medium, while the medium of the other group was supplemented with EVs at a concentration of 800 ng/mL (the maximum EV concentration isolated each time). The medium was refreshed 3 days a week for 2 more weeks.

**Table 1.** Concentrations of serum substitute medium components [19]

| Component | Concentration |
| --- | --- |
| HSA | 1% w/v |
| Anti-Anti | 1% v/v |
| ITS-G | 1% v/v |
| GlutaMax | 1% v/v |
| NEAA | 1% v/v |
| CD-lipid | 0.1% v/v |
| b-FGF | 10 ng/mL |
| rhBMP-2 | 100 ng/mL |

#### 2-5-2 ALP assay on cells

ALP is an enzyme expressed by osteoblasts and present of the cell surface. To measure the ALP activity of osteoblasts, cell-seeded wells (n=6 per group) were rinsed thoroughly with PBS several times and incubated in 250 μL of 0.2% (v/v) Triton X-100 and 5 mM MgCl_2_ solution for 30 minutes at room temperature to disturb the cell membrane. Next, the content of the well was transferred into an Eppendorf tube and centrifuged at 3000g for 10 minutes. In a 96-well plate, 80 μL of the supernatant was mixed with 20 μL of 0.75 M 2-amino-2-methyl-1-propanol buffer and 100 μL 10 mM p-nitrophenylphosphate solution and incubated until colour developed, before 0.2 M NaOH were added to stop the conversion of p-nitrophenylphosphate to p-nitrophenol. Absorbance was measured in at 405 nm on a plate reader and ALP activity was calculated by comparison to standards of known p-nitrophenol concentration.

#### 2-5-3 DNA assay

After performing ALP assay, the remaining solution was used to measure the amount of DNA per well (n=6 per group) as a measure for the number of cells. Briefly, 90 μL of the remaining solution after ALP assay was mixed with 90 μL of digestion buffer (500 mM phosphate buffer, 5 mM L-cystein (Sigma-Aldrich, C1276), 5 mM EDTA (Sigma-Aldrich, 1.08421.1000)) containing 140 μg/ml papain (Sigma-Aldrich, P-5306) at 60°C overnight in a water bath shaker (300 RPM). Next, samples were centrifuged at 3000g for 10 minutes and the DNA concentration was measured by Qubit dsDNA HS assay kit (Life Technologies, Q32851). Briefly, 10 μL of samples and standards were mixed thoroughly with DNA buffer containing DNA reagent (1:200) and the DNA concentration was measured with Qubit 2.0 Fluorometer.

#### 2-5-4 Immunohistochemistry (IHC)

Cell-seeded wells were rinsed with PBS and fixed with 10% neutral buffered formalin for 30 minutes at 4°C. To stain and detect the mineral deposition within the matrix, OsteoSense 680 EX (28422525, PerkinElmer) was used. OsteoSense 680 EX has a high affinity to hydroxyapatite both *in vitro* and *in vivo*. The dye was diluted in PBS (1:200), added to cell-seeded wells, and incubated at 37°C overnight in dark. The next day, the wells were rinsed with PBS twice and covered with 100 μL 0.5% (v/v) Triton-X 100 (Merck, 1.08603.1000) in PBS for 5 minutes to permeabilize cells. Then, the cells were rinsed with PBS and incubated with 5% (v/v) normal donkey serum and 1% (w/v) BSA (Roche, 10.7350.86001) in PBS for 1 hour at room temperature to block the non-specific antibody binding. Wells were then incubated overnight at 4°C with the primary antibody (Table 2). The wells were washed with PBS 3 times and incubated for 1 hour with the secondary antibody solution (Table 2). This was followed by 3 times rinsing of the wells with PBS, nuclei, actin cytoskeleton, and collagen were stained with 4′,6-diamidino-2-phenylindole (DAPI (Sigma-Aldrich, D9542), diluted to 0.1 μg/ml in PBS), 50 pmol Phalloidin-TRITC, and 1 μM CNA35-OG [26], respectively, for 30 minutes. Wells were rinsed with PBS 3 times and then covered with PBS. The expression of proteins was visualized with Zeis Axio Observer Z/Apotome microscope and images were processed with ImageJ (version 1.53f51). Figures were chosen to be representative images per group for all the samples assessed.

**Table 2.**
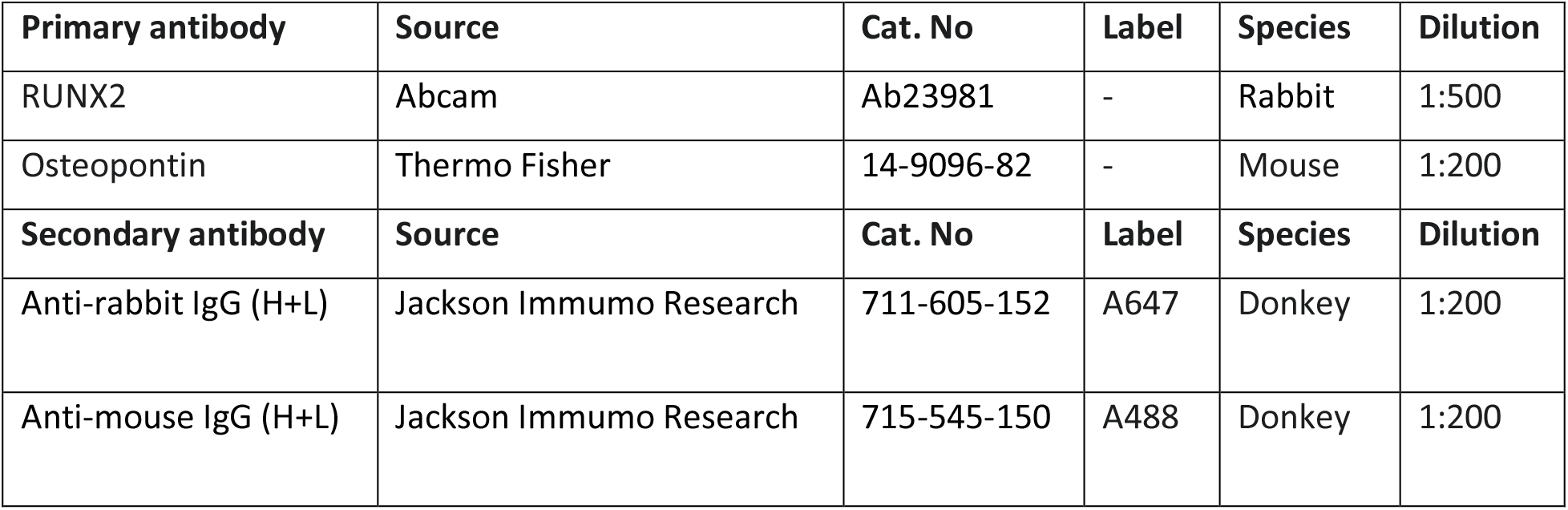
Overview of antibodies used for immunohistochemistry.

#### 2-5-5 Measure the amount of deposited phosphate within ECM

hBMSCs were cultured in serum substitute medium containing the osteogenic differentiation factors with or without the presence of EVs for 3 weeks. At the end of the culture, the deposited phosphate on the cell-seeded well plates was measured (n=4 per group). Cell-seeded wells were rinsed with PBS and incubated in 500 μL of 5% trichloroacetic acid (TCA, Sigma-Aldrich T6399) in UPW for 30 minutes. Next, the content of the each well was transferred into an Eppendorf tube and incubated for 48 hours at room temperature. Then the solids were separated by centrifuging at 3000g for 10 minutes. Eighty μL of 1:200 (v/v) diluted samples in UPW were mixed with 20 μL of working reagent and incubated for 30 minutes at room temperature. Absorbance was measured in at 620 nm on a plate reader and phosphate concentration was calculated by comparison to a phosphate standard provided in the kit.

### 2-6 Statistics

GraphPad Prism 9.0.2 (GraphPad Software, La Jolla, CA, USA) was used to perform statistical analysis and to prepare the graphs. As tested with Shapiro-Wilk test, the data were normally distributed. To test for differences, an independent t-test for particle size, mode, and concentration (Figure 2C, D, E), amount of protein (Figure 3A), ALP activity of EVs and the amount of free phosphate in the medium (Figure 6), amount of DNA and ALP activity of cells (Figure 7), and deposited phosphate (Figure 9C) was performed. The data were presented as mean +/-standard deviation and differences were considered statistically significant at a level of p-value<0.05.

**Figure 2.**
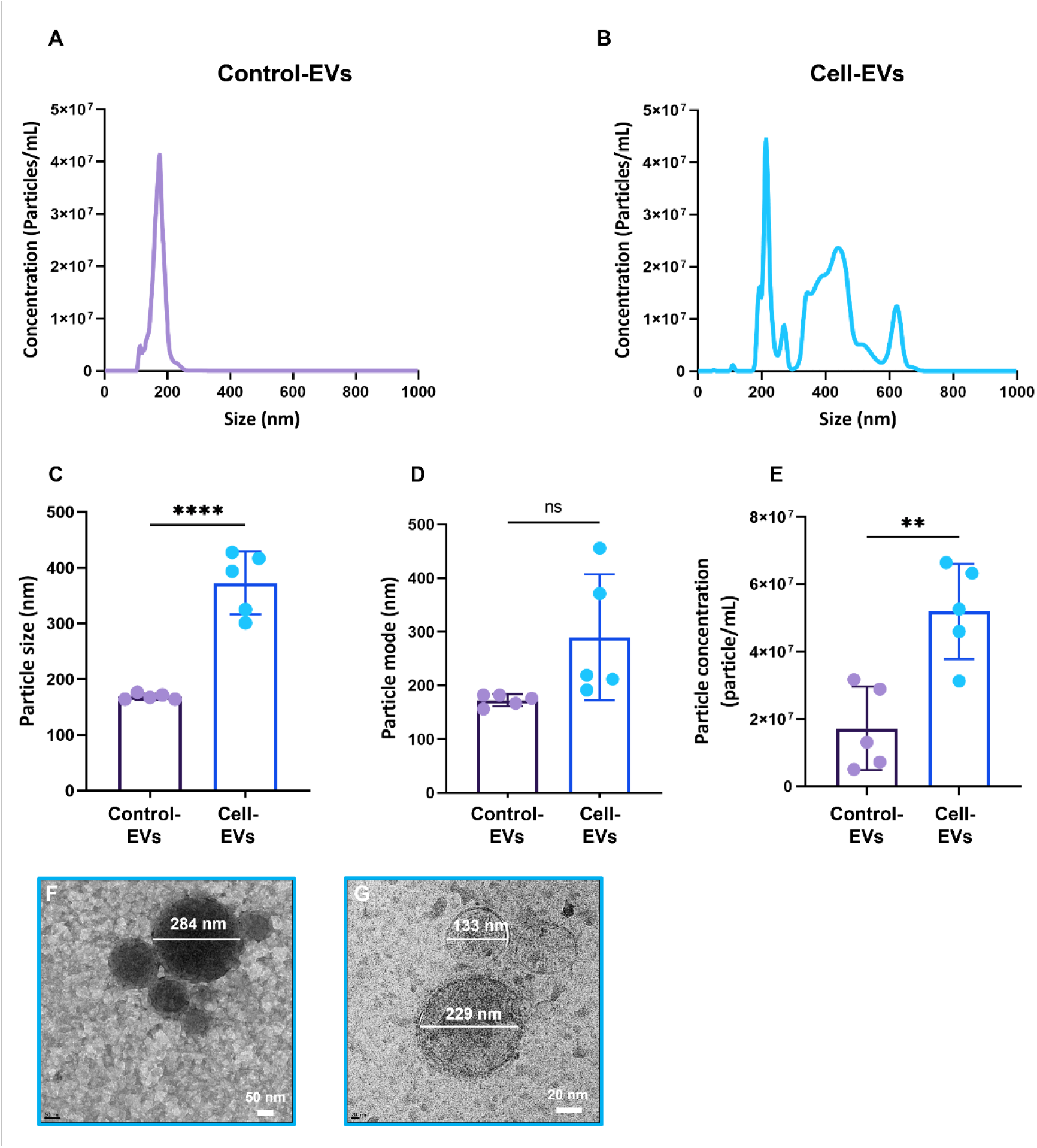
Distribution of Control-EVs (A) and Cell-EV (B) showed larger mean particle size and mode in Cell-EVs compared to Control-EVs. The particle size and mode of Cell-EVs were significantly larger than of Control-EVs (C and D). The measured particle concentration by NTA in Cell-EVs group was also significantly higher than in Control-EVs (E). Cryo-TEM images confirmed the existence of a heterogeneous population of Cell-EVs in their diameter and electron dense regions (F). (**p<0.01, ***p<0.001).

**Figure 3.**
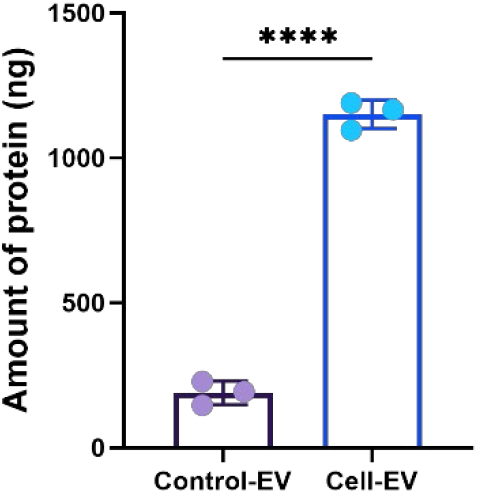
Micro BCA assay showed a significantly higher amount of protein in Cell-EV compared to Control-EV (A) (****p<0.0001).

## 3- Results

EVs were isolated from culture medium of hBMSCs differentiating towards osteoblasts/osteocytes in a previously developed 3D *in vitro* woven bone model (Cell-EVs) [16]. As a control, EVs were isolated from control medium (containing 10% FBS) without having had cell contact (Control-EVs). EVs were characterized based on their morphological, biological, and functional properties.

### 3-1 Morphological properties of EVs

NTA showed the size distribution of EVs isolated from hBMSCs during osteogenic differentiation (Cell-EVs) and control medium (Control-EVs) (Figure 2). The particle size of Cell-EVs was shown to be significantly larger and more heterogenous compared to the ones of Control-EVs with smaller and more homogenous particles (Figure 2A and B). The main peak of the Control-EVs seemed to be present in the Cell-EVs graph, but Cell-EVs graph contains more peaks that could be attributed to the EVs secreted by cells to the medium (Figure 2A and B). The measurement revealed that the mean particle size and mode (most frequent population of particle sizes) of Control-EVs were 168.9 +/-2.4 nm and 172.6 +/-5.0 nm, respectively, and the Cell-EVs had a mean particle size of 373.0 +/-25.3 nm and mode of 289.9 +/-52.4 nm (Figure 2C and D). The particle concentration measured by NTA was 5.19*10^7^ +/-6.34 *10^6^ particles/mL in Cell-EVs which was significantly higher than Control-EVs with particle concentration of 1.72*10^7^ +/-5.52*10^6^ particles/mL (Figure 2E). Cryo-TEM showed the presence of Cell-EVs with heterogeneity diameters which correlates with the NTA result of Cell-EVs (Figure 2F and G). No particles could be visualized in Control-EVs with cryo-TEM which may be due to their low concentration. The Cell-EVs were electron-dense (Figure 2F) and electron-lucent (Figure 2G) as shown by TEM images, which could suggest mineral accumulation or protein aggregation within some EVs.

### 3-2 Biological properties of EVs

The total amount of protein within Cell-EVs and Control-EVs was measured using a Micro BCA assay. The amount of protein in Cell-EVs isolated from 15 mL of cell culture supernatant was 1150.39 +/-48.79 ng which was significantly higher than the amount of protein in Control-EVs isolated from 15 mL control medium which was 189.45 +/-41.15 ng (Figure 3A). EVs were further characterized using flow cytometry. CD-9 and annexin A5, two EV surface markers were used. CD-9 is one of the general markers present on the surface of EVs [27]. Flow cytometry analysis demonstrated that Control-EVs did not show any CD-9 positive events while 1.11% of isolated Cell-EVs were CD-9 positive (Figure 4A and 5A). Annexin A5 is one of the annexin proteins that are present at substantial concentrations on the MtV membrane [28]. Flow cytometry analysis revealed that 1.58% of Control-EVs and 53.67% of Cell-EVs were annexin A5 positive (Figure 4B and 5B). This indicates the presence of MtV in the Cell-EV fraction. Double staining of the Control-EVs and Cell-EVs revealed that none of the Control-EVs were both CD-9 and annexin A5 positive while a small percentage (0.13%) of isolated Cell-EVs showed to be both CD-9 and annexin A5 positive (Figure 4C and 5C).

**Figure 4.**
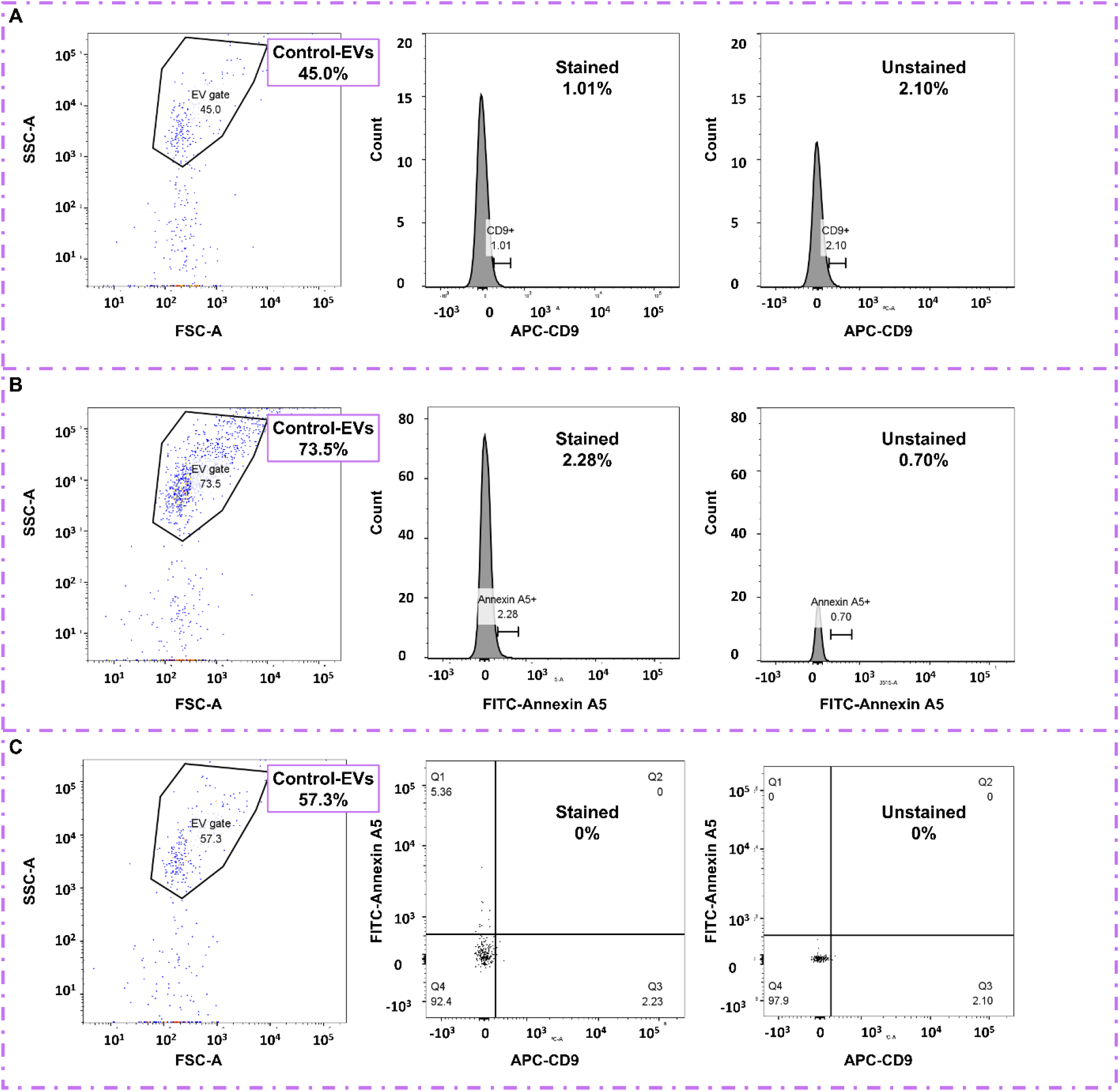
Flow cytometry analysis demonstrated that Control-EVs did not show any CD-9 positive events (A). These Control-EVs were 1.58% annexin A5 positive (B). Double staining of Control-EVs did not also show any positive events (C).

**Figure 5.**
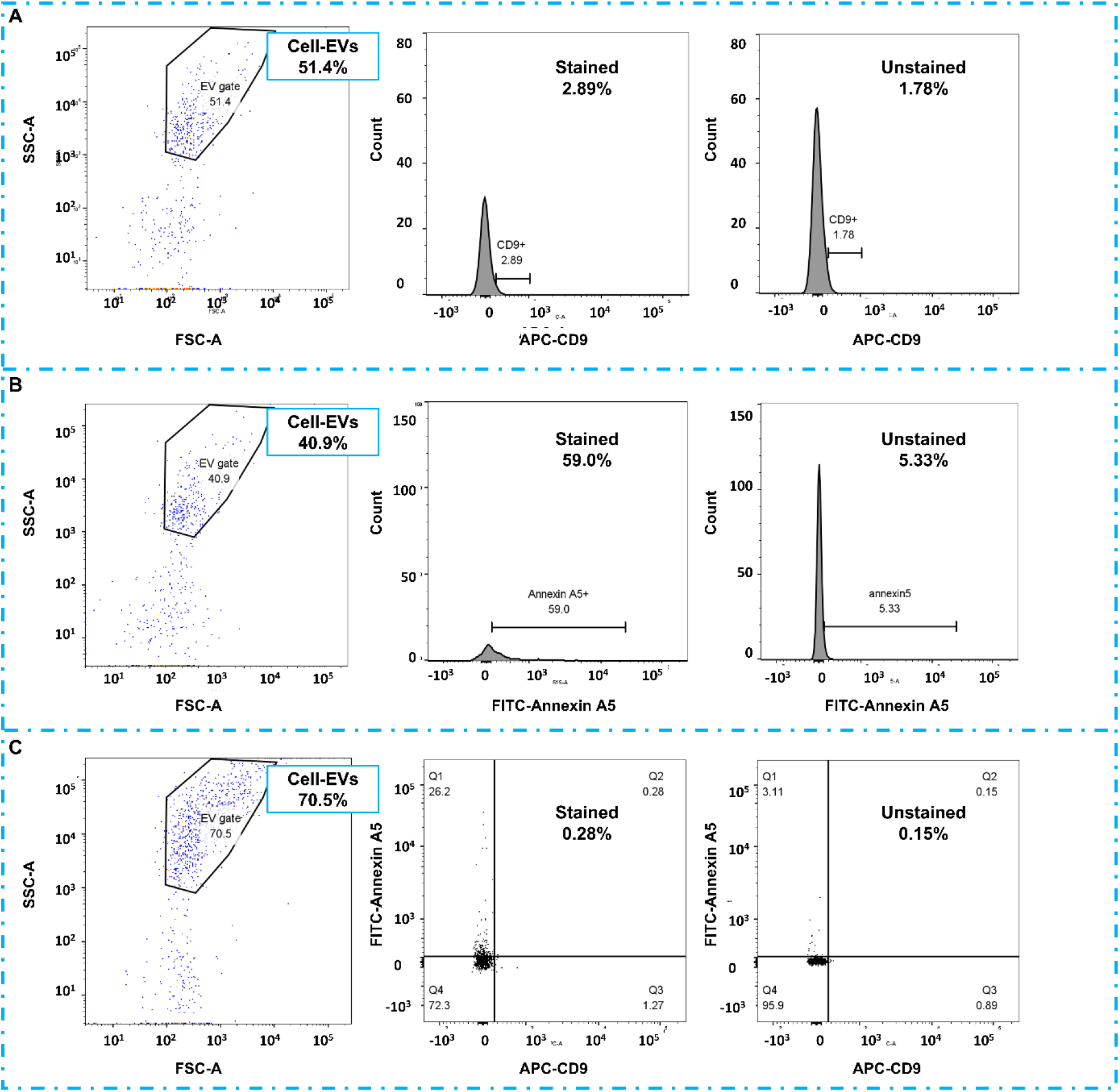
Flow cytometry analysis demonstrated that 1.11% of Cell-EVs were CD-9 positive (A). These Cell-EVs were 53.67% annexin A5 positive (B). Double staining of Control-EVs showed 0.13% positive events (C).

### 3-3 Functionality of EVs

#### 3-3-1 ALP activity of Cell-EVs and its influence on the release of free phosphate from a phosphate source

ALP, an enzyme present on the MtV membrane and responsible for dephosphorylation of pyrophosphates, was measured [29]. The ALP activity of Cell-EVs was significantly higher than the ALP activity of the same amount of Control-EVs (Figure 6A), knowing that there was lower concentration of particles in Control-EVs group. These results suggested that if Cell-EVs were incubated with a phosphate source such as β-GP, the EVs would be able to release free phosphate to the culture medium - an effect that is usually mediated by active osteoblasts. To test this, serum substitute medium containing 10 mM β-GP was incubated without or with 800 ng/mL Cell-EVs. The results indicated that the amount of free phosphate in the medium supplemented with Cell-EVs was significantly higher compared to when no Cell-EVs were added to the medium (Figure 6B). This effect confirmed the presence of active ALP on the surface of Cell-EVs.

**Figure 6.**
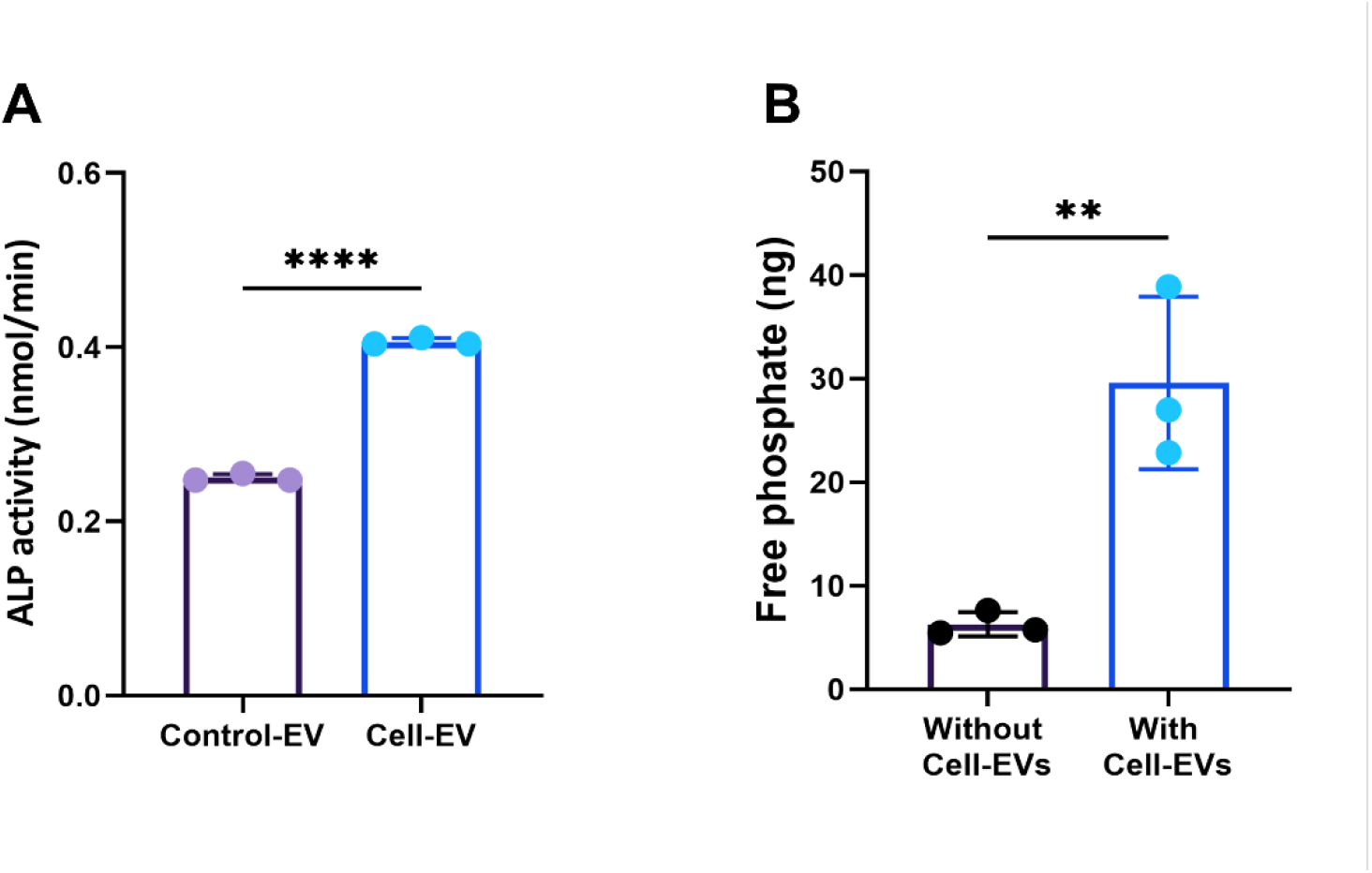
Cell-EVs showed significantly higher ALP activity compared to Control-EVs (A). The amount of free phosphate increased significantly in the medium supplemented with Cell-EVs and 10 mM β-GP compared to when no Cell-EVs were added to the medium containing 10 mM β-GP (B) (** p<0.01), ****p<0.0001).

#### 3-3-2 Influence of Cell-EVs on osteogenic differentiation of hBMSCs Cell-EVs increased the amount of DNA and ALP activity of cells

The influence of Cell-EVs on osteogenic differentiation of hBMSCs was studied by culturing cells in serum substitute medium supplemented with osteogenic differentiation factors (dexamethasone, ascorbic acid, and β-GP) and Cell-EVs. As a control, hBMSCs were cultured in serum substitute medium without the presence of Cell-EVs, but with osteogenic differentiation factors. The amount of DNA as a measure of number of cells increased significantly in the presence of Cell-EVs (Figure 7A). The addition of Cell-EVs with their own ALP activity to the cells also increased the ALP activity of the cells significantly compared to when no EVs were added to the cells (Figure 7B). Next, the measured ALP activity was normalized to the amount of DNA which also showed a significant increase in the ALP activity of cells in the presence of Cell-EVs compared to the control group (without Cell-EVs) (Figure 7C).

**Figure 7.**
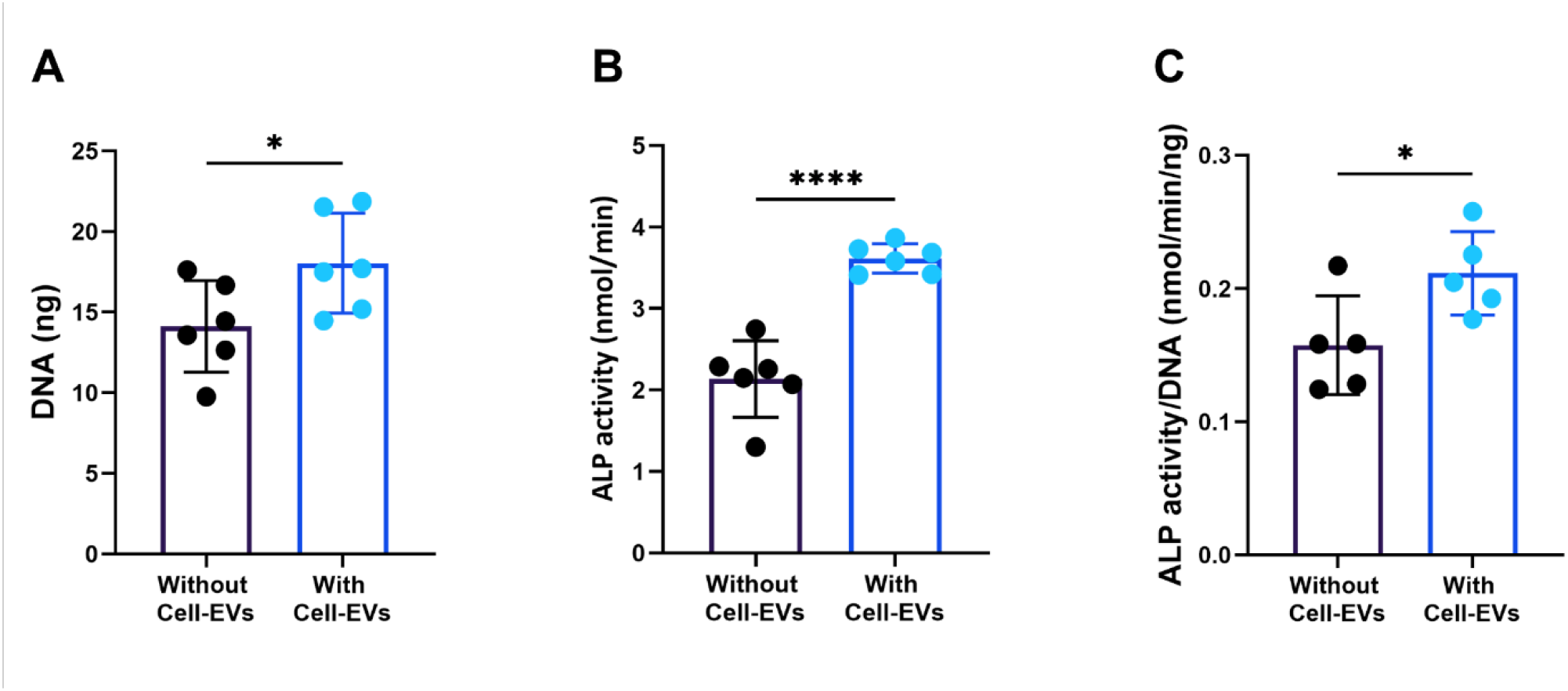
The amount of DNA (A), the ALP activity of cells with the presence of Cell-EVs increased significantly compared to when no Cell-EVs were added (B) ALP activity of cells was normalized to the DNA of cells and showed significant influence of Cell-EVs on ALP activity of cells during osteoblast differentiation (C) (* p<0.05, **** p<0.0001).

### Cell-EVs did not have an effect on the osteogenic differentiation of hBMSCs

hBMSCs were cultured in osteogenic differentiation medium supplemented with or without Cell-EVs. After 21 days of culture, osteoblast specific markers such as RUNX-2 and osteopontin were expressed in both groups (Figure 8A and B). The results revealed that adding Cell-EVs to the osteogenic culture medium did not affect cell differentiation and production of ECM proteins. Nonetheless, Cell-EVs also did not hinder osteoblastic differentiation of hBMSCs.

**Figure 8.**
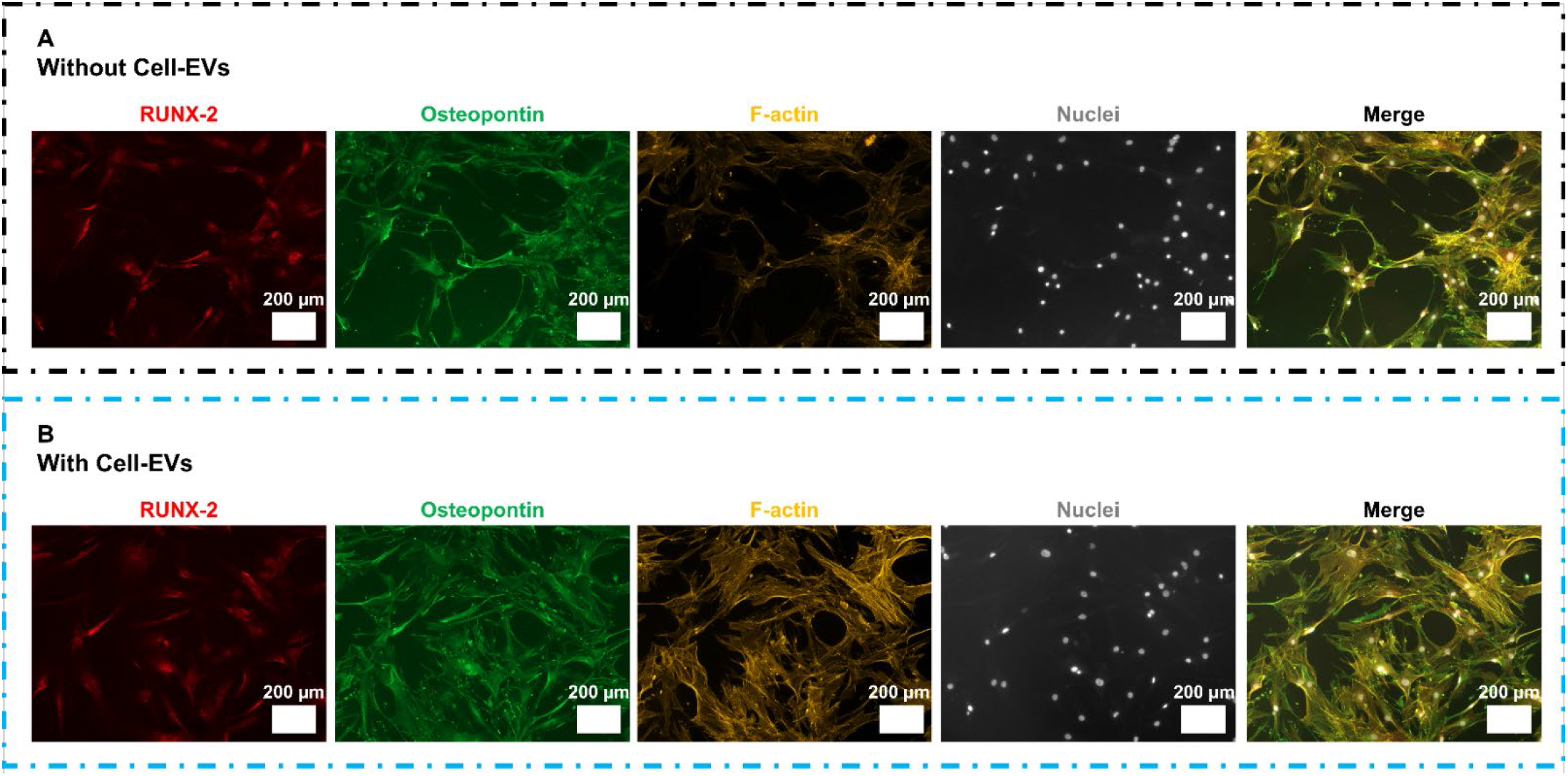
After 21 days of culturing hBMSCs in the absence (A) and presence (B) of Cell-EVs, no differences could be detected in the expression of RUNX-2 and osteopontin, and F-actin.

### Cell-EVs did not have an effect on collagen production but increased mineral deposition

Even though adding Cell-EVs to the cells cultured in osteogenic differentiation medium did not have any influence on osteoblast differentiation of hBMSCs, there seemed to be an influence on the production of ECM, particularly ECM mineralization, when Cell-EVs were added to the cells. Collagen type 1 as the main ECM protein of bone was laid down both in the presence and absence of Cell-EVs (Figure 9). The main effect of adding Cell-EVs to the medium was in mineral deposition. In the cell culture medium without Cell-EVs, only small mineral nodules were formed (Figure 9A), while there were distinctly more, and larger mineral spots present in the group where Cell-EVs were added (Figure 9B). The measurement of the phosphate in the ECM also showed significantly more deposited phosphate after 21 days in the presence of Cell-EVs compared to when no EVs were added to the medium (Figure 9C).

**Figure 9.**
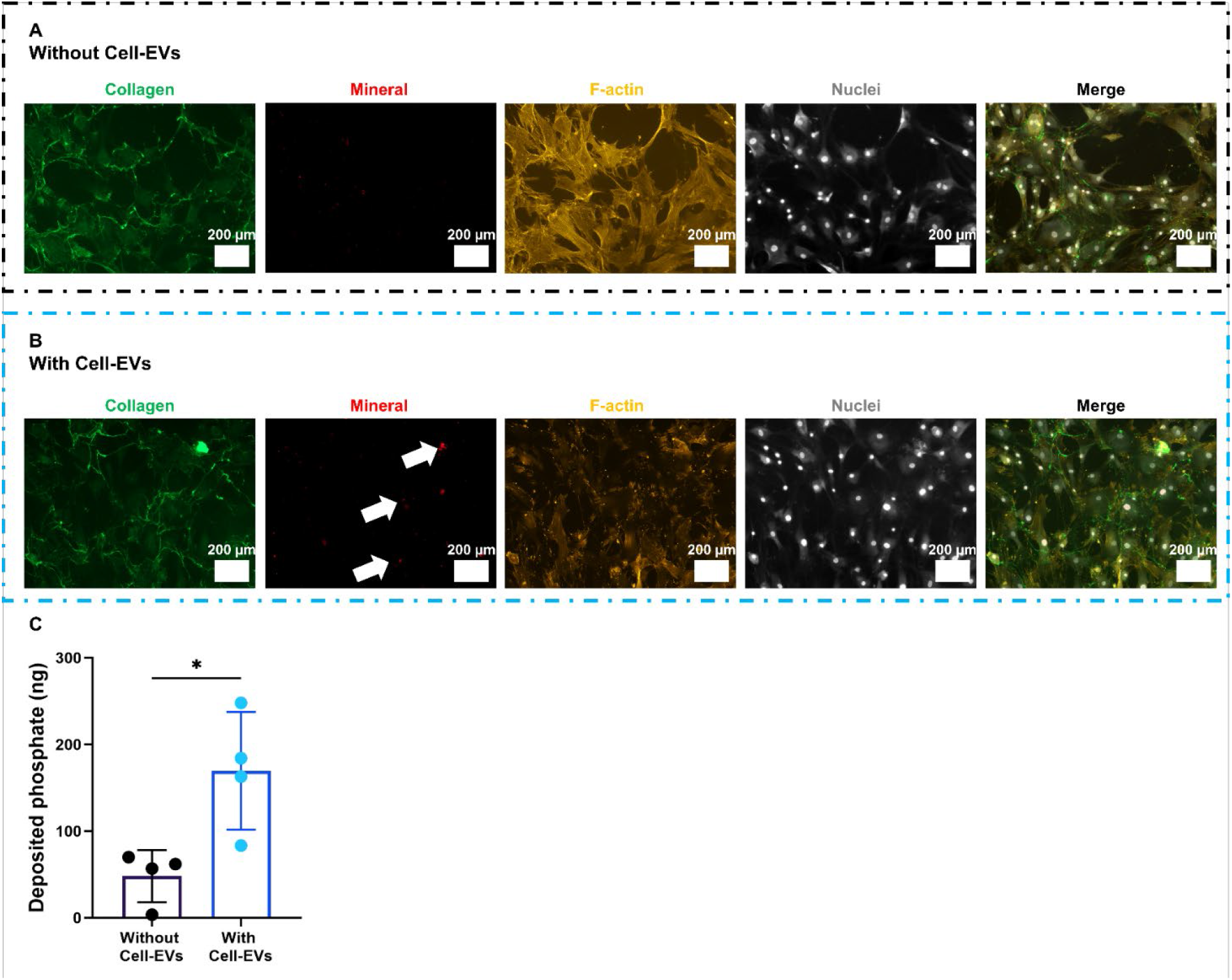
Collagen type 1 was expressed regardless of the presence of Cell-EVs in the culture (A and B). The presence of Cell-EVs in the culture increased the mineral deposition after 21 days of culture (A and B) and phosphate assay (C) (* p<0.05).

## Discussion

EVs released by osteoblasts have shown remarkable potential in osteogenesis, osteoclastogenensis, and mineralization of the organic matrix [5,7,8,30]. A subset of osteoblast-derived EVs is known as matrix vesicles (MtVs) which are involved in matrix mineralization and are equipped with mineralization-specific components such as phosphatases, calcium, and inorganic phosphate. These MtVs also showed to accelerate the osteogenic differentiation of MSCs *in vitro*, have bone-targeting potential, and induce bone formation *in vivo* [12,13]. The osteoblast-derived EVs or MtVs studies have been mostly isolated from cells cultured in two-dimensional (2D) tissue culture flasks. In our study, we investigated whether MtVs could be isolated from the cells cultured in the 3D human *in vitro* woven bone constructs, an environment with more similarity to the complex environment of *in vivo* situation [16]. Woven bone forms during skeletal development where rapid pace of matrix mineralization is needed [17]. In this process, pre-osteoblasts lay down randomly oriented collagen that becomes highly mineralized [18]. During this rapid pace mineralization process, MtVs might be highly secreted and induce the matrix mineralization. The *in vitro* woven bone construct was created through the differentiation of hBMSCs seeded on silk fibroin scaffolds and placed inside spinner flask bioreactors which induce wall shear stress to cells [16,23]. hBMSCs were differentiated into a functional 3D co-culture of osteoblasts and osteocytes and formed a mineralized matrix mimicking native woven bone [16]. In the current study, we isolated the osteoblast-derived EVs or MtVs released by cells cultured in this setup and it was hypothesized that EVs released into the culture medium show similar characteristics as MtVs, such as rich in annexin A5, exhibit membrane phosphatases such as ALP, and induce mineral deposition. EVs released by hBMSCs during osteoblast differentiation were isolated and characterized based on their morphological, biological, and functional properties. The latter was done through culturing hBMSCs in the presence and absence of EVs.

The released EVs were isolated from the cell supernatant during 4 weeks of culture. To isolate EVs, size-exclusion chromatography with columns specified for isolating EVs in the size range of 70-1000 nm has been used. This method, unlike the commonly used ultracentrifugation method, is faster, maximizes the purity of EVs, and prevents EV degradation, aggregation, and fusion due to intense gravitational force [31– 35]. The mean particle size of isolated EVs from the cell culture supernatant was 373.0 +/-25.3 nm which was at the same size range of ectosome-like MtVs [11]. Ectosome-like MtVs are formed by budding and pinching off from the membrane of mineralization-mediating cells. The membrane of such MtVs is enriched with PS-binding annexins such as annexin A5 and ALP facilitates the accumulation of calcium and phosphate ions within MtVs [36,37]. We confirmed the presence of EVs by showing the presence of the transmembrane protein CD-9, one of the EV marker proteins [27]. We also confirmed the presence of annexin A5 on the membrane of a part (not all) of Cell-EVs. The isolated EVs also contained active ALP which was able to release free phosphate from the phosphate source. These results indicated that at least a part of the isolated Cell-EVs shared similar characteristics as ectosome-like MtVs.

Cell-EVs were isolated from cell culture supernatant containing FBS. It is widely known that FBS itself contains EVs which might be co-isolated with the EVs of interest and contaminate the population of Cell-EVs [25]. To avoid EV contamination, cells have been typically cultured in EV-depleted medium or serum-free medium. However, this approach could cause cellular stress, changes in cellular phenotype, and significantly affected the growth and viability of cells [25,38–40]. In the present study, EVs were isolated from control medium (containing FBS and not in contact with cells) to investigate if Control-EVs could have been co-isolated with Cell-EVs. Although, isolated Control-EVs showed significantly less protein content and ALP activity compared to Cell-EVs, this fraction still expressed annexin A5. These results indicated that control medium contained a small population of MtVs which formed a subpopulation of the Cell-EVs, but still there were substantially more EVs present in Cell-EVs. To avoid the Control-EVs co-isolation with Cell-EVs, specialized serum substitute medium for osteogenic differentiation cultures could be used to develop an *in vitro* woven bone and isolate the secreted MtVs [19]. Using a defined medium would guarantee that only EVs expressed by cells are being isolated.

The functional properties of Cell-EVs were investigated through culturing hBMSCs in serum substitute medium supplemented with osteogenic differentiation factors and with or without Cell-EVs. Use of serum substitute medium was to prevent the influence of FBS and its EVs on cells. We previously showed that different types of FBS contains various levels of active ALP that might be due to the different levels of MtVs present in FBS [24]. These different FBS types significantly affected the matrix mineralization which was correlated with the level of active ALP in each FBS brand [24]. Our results indicated that RUNX-2 and osteopontin were expressed during osteogenic differentiation of hBMSCs regardless of the presence of Cell-EVs and these EVs did not hinder the osteogenic differentiation potential of hBMSCs, but the presence of Cell-EVs significantly increased the cellular ALP activity after 3 weeks of culture. Lack of differences in expression of osteoblast-specific factors in the presence and absence of Cell-EVs could be due to either the presence of potent osteogenic differentiation factors which could overshadow the effect of Cell-EVs or low concentration of Cell-EVs added to osteogenic differentiation medium. A previous study has also demonstrated that osteoblast-derived EVs did not induce any significant changes in osteoblastic differentiation of hBMSCs when added to medium supplemented with osteogenic differentiation factors [13]. To determine the influence of Cell-EVs, only on osteogenic differentiation of hBMSCs, addition of EVs to the medium without the presence of osteogenic differentiation factors could be studied in the future. However, the presence of β-GP, as a phosphate source in the medium, might be required. In most studies, the concentration of added EVs to the osteogenic culture medium was reported to be between 2.5 to 10 μg/mL [12,13,41]. The highest Cell-EV concentration that we could isolate from the cell culture medium collected during 4 weeks of experiment using size-exclusion chromatography was 800 ng/mL. Since MtVs can also bind to collagen matrix, for example through annexin proteins and ALP, the use of collagenase before isolation of osteoblast-derived EVs from osseous tissue was reported as an approach to release MtVs from the matrix [42]. This approach resulted in a population of MtVs with a higher ALP activity and mineral formation potential compared to the non-collagenase treated osseous tissue [43]. A potential way to increase the concentration of isolated Cell-EVs could be breaking down the collagen matrix using collagenase [44,45]. In this case, the effect of collagenase on the characteristics of Cell-EVs should be investigated as well.

Matrix mineralization is an important step in *in vitro* bone tissue formation. MtVs are known to play an essential role in mineralization of the organic bone matrix. These vesicles can bind to the collagen matrix. The membrane proteins of MtVs facilitate entry of calcium and phosphate ions into MtVs. These ions then form amorphous or crystalized minerals inside the MtVs which can rupture the membrane and form mineral nodules in the ECM [10,11]. Adding Cell-EVs to hBMSCs showed significant increase in deposited phosphate compared to when hBMSCs were continued to be cultured in serum substitute medium in the absence of Cell-EVs. The microscopy images also confirmed the formation of mineral nodules on the deposited collagenous matrix. The mineral formation could be directly due to the contribution of ALP activity of Cell-EVs on increasing the amount of free phosphate in the medium or indirectly through the increase in cellular ALP activity in the presence of Cell-EVs. Previous studies also demonstrated the influence of MtVs on increasing the cellular ALP activity and matrix mineralization [12,13,41].

Previous studies have shown that EVs isolated from different stages of osteogenic differentiation of MSCs showed different characteristics [46–48]. A recent study revealed that EVs isolated on days 21, 28, and 35 of osteogenic differentiation of hBMSCs have different potentials in hBMSCs proliferation, osteoblast differentiation, and ECM mineralization [46]. It has been shown that EVs isolated from earlier time points exhibited a high proliferation stimulus; while EVs isolated from later time points resulted in more osteogenic differentiation and mineral deposition [46]. In our study, EVs were isolated from the mixture of osteogenic medium collected during 28 days of culture. For future studies, to increase the purity of MtVs, either EVs secreted by hBSMCs during later stages of osteogenic differentiation using the same setup could be isolated or another step of isolation could be added to the current isolation method. For instance, after isolating EVs using size-exclusion chromatography, the affinity chromatography method could be used to isolate EVs with specific surface markers, for example, ALP [49–51].

Osteoblast derived EVs and more specifically MtVs are increasingly being considered as a promising therapeutic agent for bone regeneration and fracture healing [7,12,15]. Considering advances in the development of *in vitro* bone models using bone tissue engineering approaches [52], more physiologically relevant models such as our previously developed 3D human *in vitro* woven bone constructs could be used to guide MtV-producing cells to produce targeted MtVs *in vitro*. Later, these secreted MtVs could be integrated in biomaterials and delivered at a controlled release rate to the site of interest to regulate bone regeneration locally. The future studies should focus on development of such MtV-integrated biomaterials for therapeutic applications for bone tissue.

## Conclusion

We have shown that our previously developed 3D human *in vitro* woven bone models could result in secretion of MtVs from osteogenic differentiated hBSMCs. Secreted EVs during osteogenic differentiation of hBSMCs were isolated using size-exclusion chromatography and characterized based on their morphological, biological, and functional properties. A part of these EVs shared the similarities as MtVs in size, CD-9 and Annexin A5 surface markers, ALP activity, increase the concentration of free phosphate in the medium as well as deposited phosphate on ECM. These isolated MtVs could potentially be used as a biological agent for bone regeneration and fracture healing.

## Conflict of interest

The authors declare that there is no conflict of interest.

## Acknowledgement

This work has been financially supported by the Dutch Ministry of Education, Culture and Science (Gravitation Program 024.003.013). We would like to thank Rick Joosten for his help in preparing and imaging TEM samples.

## Notes

### Competing Interest Statement

The authors have declared no competing interest.

## References

[1] Clarke B. Normal bone anatomy and physiology. Clin J Am Soc Nephrol 2008;3 Suppl 3:131–9. https://doi.org/10.2215/CJN.04151206.

[2] Caplan AI. Mesenchymal stem cells. J Orthop Res 1991;9:641–50. https://doi.org/https://doi.org/10.1002/jor.1100090504.

[3] Reznikov N, Bilton M, Lari L, Stevens MM, Kröger R. Fractal-like hierarchical organization of bone begins at the nanoscale. Science (80-) 2018;360. https://doi.org/10.1126/science.aao2189.

[4] Hart NH, Nimphius S, Rantalainen T, Ireland A, Siafarikas A, Newton RU. Mechanical basis of bone strength: Influence of bone material, bone structure and muscle action. J Musculoskelet Neuronal Interact 2017;17:114–39. https://doi.org/10.5586/asbp.2013.028.

[5] Tao S-C, Guo S-C. Extracellular vesicles in bone: “dogrobbers” in the “eternal battle field.” Cell Commun Signal 2019;17:6. https://doi.org/10.1186/s12964-019-0319-5.

[6] Boonrungsiman S, Gentleman E, Carzaniga R, Evans ND, McComb DW, Porter AE, et al. The role of intracellular calcium phosphate in osteoblast-mediated bone apatite formation. Proc Natl Acad Sci 2012;109:14170–5. https://doi.org/10.1073/pnas.1208916109.

[7] Azoidis I, Cox SC, Davies OG. The role of extracellular vesicles in biomineralisation: current perspective and application in regenerative medicine. J Tissue Eng 2018;9:2041731418810130. https://doi.org/10.1177/2041731418810130.

[8] Kobayashi-Sun J, Yamamori S, Kondo M, Kuroda J, Ikegame M, Suzuki N, et al. Uptake of osteoblast-derived extracellular vesicles promotes the differentiation of osteoclasts in the zebrafish scale. Commun Biol 2020;3:1–12. https://doi.org/10.1038/s42003-020-0925-1.

[9] Uenaka M, Yamashita E, Kikuta J, Morimoto A, Ao T, Mizuno H, et al. Osteoblast-derived vesicles induce a switch from bone-formation to bone-resorption in vivo. Nat Commun 2022;13:1066. https://doi.org/10.1038/s41467-022-28673-2.

[10] Hasegawa T, Hongo H, Yamamoto T, Abe M, Yoshino H, Haraguchi-Kitakamae M, et al. Matrix Vesicle-Mediated Mineralization and Osteocytic Regulation of Bone Mineralization. Int J Mol Sci 2022;2022. https://doi.org/10.3390/ijms23179941.

[11] Ansari S, de Wildt BWM, Vis MAM, de Korte CE, Ito K, Hofmann S, et al. Matrix Vesicles: Role in Bone Mineralization and Potential Use as Therapeutics. Pharmaceuticals (Basel) 2021;2021. https://doi.org/10.3390/ph14040289.

[12] Wei Y, Tang C, Zhang J, Li Z, Zhang X, Miron RJ, et al. Extracellular vesicles derived from the mid-to-late stage of osteoblast differentiation markedly enhance osteogenesis in vitro and in vivo. Biochem Biophys Res Commun 2019;514:252–8. https://doi.org/https://doi.org/10.1016/j.bbrc.2019.04.029.

[13] Davies OG, Cox SC, Williams RL, Tsaroucha D, Dorrepaal RM, Lewis MP, et al. Annexin-enriched osteoblast-derived vesicles act as an extracellular site of mineral nucleation within developing stem cell cultures. Sci Rep 2017;7:12639. https://doi.org/10.1038/s41598-017-13027-6.

[14] Emami A, Talaei-Khozani T, Tavanafar S, Zareifard N, Azarpira N, Vojdani Z. Synergic effects of decellularized bone matrix, hydroxyapatite, and extracellular vesicles on repairing of the rabbit mandibular bone defect model. J Transl Med 2020;18:361. https://doi.org/10.1186/s12967-020-02525-3.

[15] Cappariello A, Loftus A, Muraca M, Maurizi A, Rucci N, Teti A. Osteoblast-Derived Extracellular Vesicles Are Biological Tools for the Delivery of Active Molecules to Bone. J Bone Miner Res Off J Am Soc Bone Miner Res 2018;33:517–33. https://doi.org/10.1002/jbmr.3332.

[16] Akiva A, Melke J, Ansari S, Liv N, van der Meijden R, van Erp M, et al. An Organoid for Woven Bone. Adv Funct Mater 2021;n/a:2010524. https://doi.org/https://doi.org/10.1002/adfm.202010524.

[17] Uthgenannt BA, Kramer MH, Hwu JA, Wopenka B, Silva MJ. Skeletal self-repair: stress fracture healing by rapid formation and densification of woven bone. J Bone Miner Res 2007;22:1548–56. https://doi.org/10.1359/jbmr.0070614.

[18] McBride SH, Silva MJ. Adaptive and Injury Response of Bone to Mechanical Loading. Bonekey Osteovision 2012;2012. https://doi.org/10.1038/bonekey.2012.192.

[19] Ansari S, Ito K, Hofmann S. Development of serum substitute medium for bone tissue engineering. BioRxiv 2022:2022.10.07.511271. https://doi.org/10.1101/2022.10.07.511271.

[20] Tsukada M, Gotoh Y, Nagura M, Minoura N, Kasai N, Freddi G. Structural changes of silk fibroin membranes induced by immersion in methanol aqueous solutions. J Polym Sci Part B Polym Phys 1994;32:961–8. https://doi.org/10.1002/polb.1994.090320519.

[21] Hofmann S, Hagenmüller H, Koch AM, Müller R, Vunjak-Novakovic G, Kaplan DL, et al. Control of in vitro tissue-engineered bone-like structures using human mesenchymal stem cells and porous silk scaffolds. Biomaterials 2007;28:1152–62. https://doi.org/https://doi.org/10.1016/j.biomaterials.2006.10.019.

[22] Melke J, Zhao F, Ito K, Hofmann S. Orbital seeding of mesenchymal stromal cells increases osteogenic differentiation and bone-like tissue formation. J Orthop Res 2020;38:1228–37. https://doi.org/10.1002/jor.24583.

[23] Melke J, Zhao F, Rietbergen B, Ito K, Hofmann S. Localisation of mineralised tissue in a complex spinner flask environment correlates with predicted wall shear stress level localisation. Eur Cells Mater 2018;36:57–68. https://doi.org/10.22203/eCM.v036a05.

[24] Ansari S, Ito K, Hofmann S. Alkaline Phosphatase Activity of Serum Affects Osteogenic Differentiation Cultures. ACS Omega 2022;7:12724–33. https://doi.org/10.1021/acsomega.1c07225.

[25] Lehrich BM, Liang Y, Fiandaca MS. Foetal bovine serum influence on in vitro extracellular vesicle analyses. J Extracell Vesicles 2021;10:e12061. https://doi.org/10.1002/jev2.12061.

[26] Krahn KN, Bouten CVC, van Tuijl S, van Zandvoort MAMJ, Merkx M. Fluorescently labeled collagen binding proteins allow specific visualization of collagen in tissues and live cell culture. Anal Biochem 2006;350:177–85. https://doi.org/10.1016/j.ab.2006.01.013.

[27] Yoshioka Y, Konishi Y, Kosaka N, Katsuda T, Kato T, Ochiya T. Comparative marker analysis of extracellular vesicles in different human cancer types. J Extracell Vesicles 2013;2013. https://doi.org/10.3402/jev.v2i0.20424.

[28] Genge BR, Wu LNY, Wuthier RE. In vitro modeling of matrix vesicle nucleation: synergistic stimulation of mineral formation by annexin A5 and phosphatidylserine. J Biol Chem 2007;282:26035–45. https://doi.org/10.1074/jbc.M701057200.

[29] Golub EE, Boesze-Battaglia K. The role of alkaline phosphatase in Mineralization. Curr Opin Orthop 2007;18:444–8. https://doi.org/10.1084/jem.93.5.415.

[30] Deng L, Peng Y, Jiang Y, Wu Y, Ding Y, Wang Y, et al. Imipramine protects against bone loss by inhibition of osteoblast-derived microvesicles. Int J Mol Sci 2017;18:1–14. https://doi.org/10.3390/ijms18051013.

[31] Linares R, Tan S, Gounou C, Arraud N, Brisson AR. High-speed centrifugation induces aggregation of extracellular vesicles. J Extracell Vesicles 2015;4:29509. https://doi.org/10.3402/jev.v4.29509.

[32] Mol EA, Goumans M-J, Doevendans PA, Sluijter JPG, Vader P. Higher functionality of extracellular vesicles isolated using size-exclusion chromatography compared to ultracentrifugation. Nanomedicine Nanotechnology, Biol Med 2017;13:2061–5. https://doi.org/https://doi.org/10.1016/j.nano.2017.03.011.

[33] Takov K, Yellon DM, Davidson SM. Comparison of small extracellular vesicles isolated from plasma by ultracentrifugation or size-exclusion chromatography: yield, purity and functional potential. J Extracell Vesicles 2019;8:1560809. https://doi.org/10.1080/20013078.2018.1560809.

[34] Liangsupree T, Multia E, Riekkola M-L. Modern isolation and separation techniques for extracellular vesicles. J Chromatogr A 2021;1636:461773. https://doi.org/https://doi.org/10.1016/j.chroma.2020.461773.

[35] Gardiner C, Di Vizio D, Sahoo S, Théry C, Witwer KW, Wauben M, et al. Techniques used for the isolation and characterization of extracellular vesicles: results of a worldwide survey. J Extracell Vesicles 2016;5:32945. https://doi.org/10.3402/jev.v5.32945.

[36] Wuthier RE, Lipscomb GF. Matrix vesicles: structure, composition, formation and function in calcification. Front Biosci (Landmark Ed 2011;16:2812–902. https://doi.org/10.2741/3887.

[37] Kirsch T, Nah HD, Demuth DR, Harrison G, Golub EE, Adams SL, et al. Annexin V-mediated calcium flux across membranes is dependent on the lipid composition: implications for cartilage mineralization. Biochemistry 1997;36:3359–67. https://doi.org/10.1021/bi9626867.

[38] Lehrich BM, Liang Y, Khosravi P, Federoff HJ, Fiandaca MS. Fetal Bovine Serum-Derived Extracellular Vesicles Persist within Vesicle-Depleted Culture Media. Int J Mol Sci 2018;2018. https://doi.org/10.3390/ijms19113538.

[39] Aswad H, Jalabert A, Rome S. Depleting extracellular vesicles from fetal bovine serum alters proliferation and differentiation of skeletal muscle cells in vitro. BMC Biotechnol 2016;16:32. https://doi.org/10.1186/s12896-016-0262-0.

[40] Eitan E, Zhang S, Witwer KW, Mattson MP. Extracellular vesicle-depleted fetal bovine and human sera have reduced capacity to support cell growth. J Extracell Vesicles 2015;4:26373. https://doi.org/10.3402/jev.v4.26373.

[41] Qin Y, Wang L, Gao Z, Chen G, Zhang C. Bone marrow stromal/stem cell-derived extracellular vesicles regulate osteoblast activity and differentiation in vitro and promote bone regeneration in vivo. Sci Rep 2016;6:21961. https://doi.org/10.1038/srep21961.

[42] Wu LN, Genge BR, Lloyd GC, Wuthier RE. Collagen-binding proteins in collagenase-released matrix vesicles from cartilage. Interaction between matrix vesicle proteins and different types of collagen. J Biol Chem 1991;266:1195–203.

[43] Balcerzak M, Radisson J, Azzar G, Farlay D, Boivin G, Pikula S, et al. A comparative analysis of strategies for isolation of matrix vesicles. Anal Biochem 2007;361:176–82. https://doi.org/https://doi.org/10.1016/j.ab.2006.10.001.

[44] Ali SY, Sajdera SW, Anderson HC. Isolation and characterization of calcifying matrix vesicles from epiphyseal cartilage. Proc Natl Acad Sci U S A 1970;67:1513–20. https://doi.org/10.1073/pnas.67.3.1513.

[45] Nahar NN, Missana LR, Garimella R, Tague SE, Anderson HC. Matrix vesicles are carriers of bone morphogenetic proteins (BMPs), vascular endothelial growth factor (VEGF), and noncollagenous matrix proteins. J Bone Miner Metab 2008;26:514–9. https://doi.org/10.1007/s00774-008-0859-z.

[46] Wang C, Stöckl S, Li S, Herrmann M, Lukas C, Reinders Y, et al. Effects of Extracellular Vesicles from Osteogenic Differentiated Human BMSCs on Osteogenic and Adipogenic Differentiation Capacity of Naïve Human BMSCs. Cells 2022;2022. https://doi.org/10.3390/cells11162491.

[47] Wang X, Omar O, Vazirisani F, Thomsen P, Ekström K. Mesenchymal stem cell-derived exosomes have altered microRNA profiles and induce osteogenic differentiation depending on the stage of differentiation. PLoS One 2018;13:e0193059.

[48] Wei F, Li Z, Crawford R, Xiao Y, Zhou Y. Immunoregulatory role of exosomes derived from differentiating mesenchymal stromal cells on inflammation and osteogenesis. J Tissue Eng Regen Med 2019;13:1978–91. https://doi.org/https://doi.org/10.1002/term.2947.

[49] Brenna O, Perrella M, Pace M, Pietta PG. Affinity-chromatography purification of alkaline phosphatase from calf intestine. Biochem J 1975;151:291–6. https://doi.org/10.1042/bj1510291.

[50] Landt M, Boltz SC, Butler LG. Alkaline phosphatase: affinity chromatography and inhibition by phosphonic acids. Biochemistry 1978;17:915–9. https://doi.org/10.1021/bi00598a027.

[51] Latner AL, Hodson AW. Human liver alkaline phosphatase purified by affinity chromatography, ultracentrifugation and polyacrylamide-gel electrophoresis. Biochem J 1976;159:697–705. https://doi.org/10.1042/bj1590697.

[52] de Wildt BWM, Ansari S, Sommerdijk Najm, Ito K, Akiva A, Hofmann S. From bone regeneration to three-dimensional in vitro models: tissue engineering of organized bone extracellular matrix. Curr Opin Biomed Eng 2019;10:107–15. https://doi.org/10.1016/j.cobme.2019.05.005.

